# Functional analysis of a hypomorphic allele shows that MMP14 catalytic activity is the prime determinant of the Winchester syndrome phenotype

**DOI:** 10.1101/281485

**Authors:** Ivo J.H.M. de Vos, Evelyn Yaqiong Tao, Sheena Li Ming Ong, Julian L. Goggi, Thomas Scerri, Gabrielle R. Wilson, Chernis Guai Mun Low, Arnette Shi Wei Wong, Dominic Grussu, Alexander P.A. Stegmann, Michel van Geel, Renske Janssen, David J. Amor, Melanie Bahlo, Norris R. Dunn, Thomas J. Carney, Paul J. Lockhart, Barry J. Coull, Maurice A.M. van Steensel

## Abstract

Winchester syndrome (WS, MIM #277950) is an extremely rare autosomal recessive skeletal dysplasia characterized by progressive joint destruction and osteolysis. To date, only one missense mutation in *MMP14*, encoding the membrane-bound matrix metalloprotease 14, has been reported in WS patients. Here, we report a novel hypomorphic MMP14 p.Arg111His (R111H) allele, associated with a mitigated form of WS. Functional analysis demonstrated that this mutation, in contrast to previously reported human and murine *MMP14* mutations, does not affect MMP14’s transport to the cell membrane. Instead, it partially impairs MMP14’s proteolytic activity. This residual activity likely accounts for the mitigated phenotype observed in our patients. Based on our observations as well as previously published data, we hypothesize that MMP14’s catalytic activity is the prime determinant of disease severity. Given the limitations of our *in vitro* assays in addressing the consequences of MMP14 dysfunction, we generated a novel *mmp14a/b* knockout zebrafish model. The fish accurately reflected key aspects of the WS phenotype including craniofacial malformations, kyphosis, short-stature and reduced bone density due to defective collagen remodeling. Notably, the zebrafish model will be a valuable tool for developing novel therapeutic approaches to a devastating bone disorder.

## Introduction

In 2007, we reported two Dutch brothers with an autosomal recessive disorder consisting of dysmorphic facial features, mitral valve prolapse, severe acne and reduced bone density [1]. We diagnosed them with Borrone syndrome, as their phenotype strongly resembled that of two patients first described by Borrone et al. in 1993 [2]. Symptoms in our patients were less severe, which we attributed to their younger age compared to Borrone’s patients at the time of diagnosis. However, in the intervening years since diagnosis their phenotype has not appreciably worsened, suggesting that it is intrinsically milder.

In the patients originally reported by Borrone et al., we recently identified a homozygous splice site mutation in *SH3PXD2B* [3]. Thus, Borrone syndrome is no longer considered as a separate entity, but as allelic to Frank-Ter Haar syndrome (FTHS, MIM #249420) [4]. SH3 and Phox-homology (PX) Domain-containing Protein 2B (SH3PXD2B, also known as TKS4) is an adapter protein required for functionality of podosomes [5]. These are actin-rich membrane structures that mediate adhesion and invasive motility in a variety of cell types. Specifically, upon phosphorylation by c-SRC, SH3PXD2B recruits the membrane-bound matrix metalloprotease 14 (MMP14, also known as MT1-MMP) to the nascent podosome membrane [6]. Here, MMP14 hydrolyzes intact fibrillar collagen and activates downstream effectors, including the gelatinase MMP2 that in turn can further degrade fragmented collagen fibrils [7–9]. MMP14’s collagenolytic activity is thought to be one of its most important functions *in vivo*, for which homodimerization is required [10]. Loss of either MMP2 or MMP14 results in a spectrum of recessive skeletal dysplasias with osteolysis, encompassing multicentric osteolysis, nodulosis and arthropathy (MONA, MIM #259600) and Winchester syndrome (WS, MIM #277950). These disorders exhibit significant clinical overlap. Notably, WS is associated with mutations in *MMP2* as well as in *MMP14* [11, 12].

In our patients, we found no mutation or deletion of *SH3PXD2B*. Subsequent homozygosity mapping and next-generation sequencing identified a novel homozygous missense mutation in *MMP14*, c.332G>A (www.LOVD.nl/MMP14), which was predicted to result in the substitution p.Arg111His (R111H, see supplemental methods and Fig. S1) and assessed as possibly damaging by multiple algorithms (data not shown). Accordingly, we revised our patients’ diagnosis to WS, pending confirmation of the mutation’s pathogenicity.

*MMP14* encodes a membrane-bound metalloprotease that requires removal of an N-terminal pro-domain sequence for its activation and presentation at the cell surface [13]. The pro-domain has two furin cleavage motifs, R^89^-R-P-R-C^93^ and R^108^-R-K-R-Y^112^. Previously published work suggests that the latter motif is cleaved to generate the active enzyme [13, 14]. Therefore, we reasoned that the R111H mutation might interfere with cleavage and thereby impair MMP14 membrane localization and activation.

To test our hypothesis, we analyzed the consequences of the R111H change for MMP14’s intracellular processing and functionality, comparing with known mutations associated with WS and similar mouse phenotypes. To better understand the connection between loss of MMP14 activity and the clinical manifestations of WS, we additionally generated a knockout zebrafish model. Our findings provide novel insights into the pathogenesis of the WS phenotype, with potential consequences for therapy.

## Results

### An in vitro model for assessing MMP14 processing and subcellular localization

To examine MMP14 processing, we created a construct encoding either wild type (WT) or mutant human pro-MMP14 with an N-terminal triple (3)-HA tag and a C-terminal EGFP (resulting in the fusion protein 3HA–MMP14–EGFP, Fig. 1A). Given correct processing of MMP14, the 3HA tag should not be detectable in a similar location to EGFP. The EGFP signal, on the other hand, should be visible at the Golgi/trans-Golgi network, the cell membrane and along the route by which the protein traffics. Aberrant processing and subsequent abnormal trafficking ought to be reflected in an altered subcellular distribution of both tags. Fusion proteins were exogenously expressed in MRC-5V1 immortalized human fetal lung fibroblasts (hereafter termed MRC5) and compared to control of 3HA-EGFP (Fig. 1B). This cell type is a good model for the disease phenotype, which includes connective tissue and lung abnormalities in humans and mouse models respectively [15, 16]. Immunoblotting with a commercially available antibody demonstrated that MRC5 cells have very low endogenous MMP14 expression (Fig. S2). Thus, mutant protein localization is unlikely to be affected by dimerization with endogenous MMP14. Next, we determined that the localization of exogenously expressed double-tagged MMP14-WT was very similar from that of endogenous MMP14 (Fig. S3). Having established that our tagging strategy did not affect protein localization, we next compared and contrasted WT MMP14 and MMP14-R111H with previously characterized MMP14 missense mutants to determine why our patients’ phenotype was comparatively mild.

**Figure 1.**
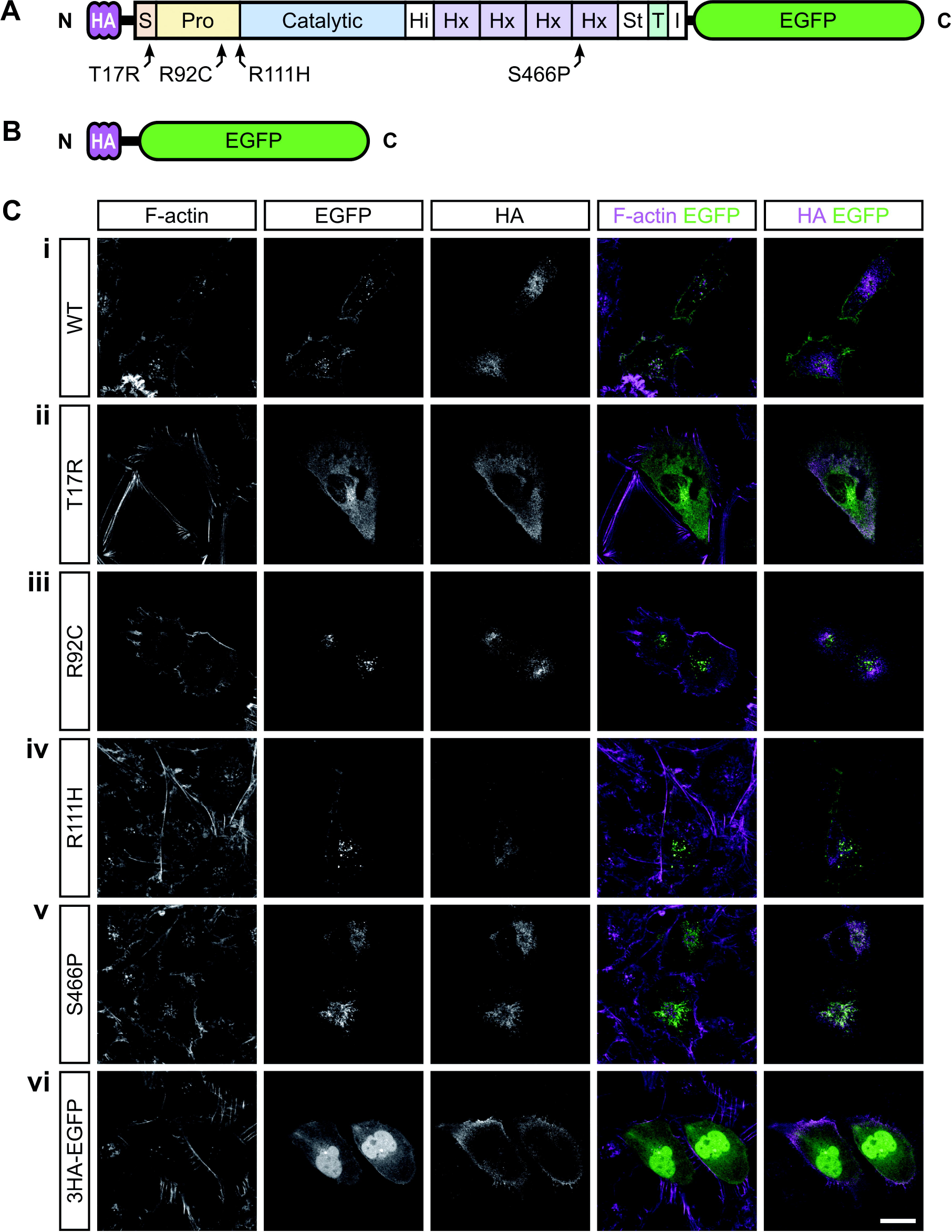
The R111H mutation does not impair MMP14 cell surface localization, in contrast to mutations T17R, R92C and S466P. **A**, schematic representation of the MMP14 fusion protein with N-terminal triple (3)-HA tag and C-terminal EGFP. Both tags are attached to MMP14 by flexible linkers (horizontal black lines). Indicated are the mutations relative to the protein domains. **B**, schematic representation of the 3HA-EGFP fusion protein which served as control. **C**, subcellular localization of WT and mutant 3HA-MMP14-EGFP fusion proteins exogenously expressed in MRC5 cells. MMP14 WT-EGFP (i) and MMP14 R111H-EGFP (iv) are present in the perinuclear region and at the cell surface, whereas the HA-tag is absent at the cell surface. For other mutant fusion proteins, cell surface localization is impaired and the two tags partially colocalize in the perinuclear region. Scale bar equals 20 μm.

### MMP14 R111H is processed normally and is trafficked to the cell surface

As shown in Figures 1C and S4, MMP14-R111H-EGFP (panel iv) localizes in a similar manner to that observed for MMP14-WT-EGFP (panel i). The lack of any HA signal at the cell membrane suggest that R111H does not markedly impact removal of the signal peptide. In Western blot (WB) analysis (Fig. 2A lane 2 (WT) and lane 5 (R111H)) the banding pattern of R111H was similar to that of WT, with EGFP signal indicating removal of the HA tag and subsequent further processing of the resultant protein.

**Figure 2.**
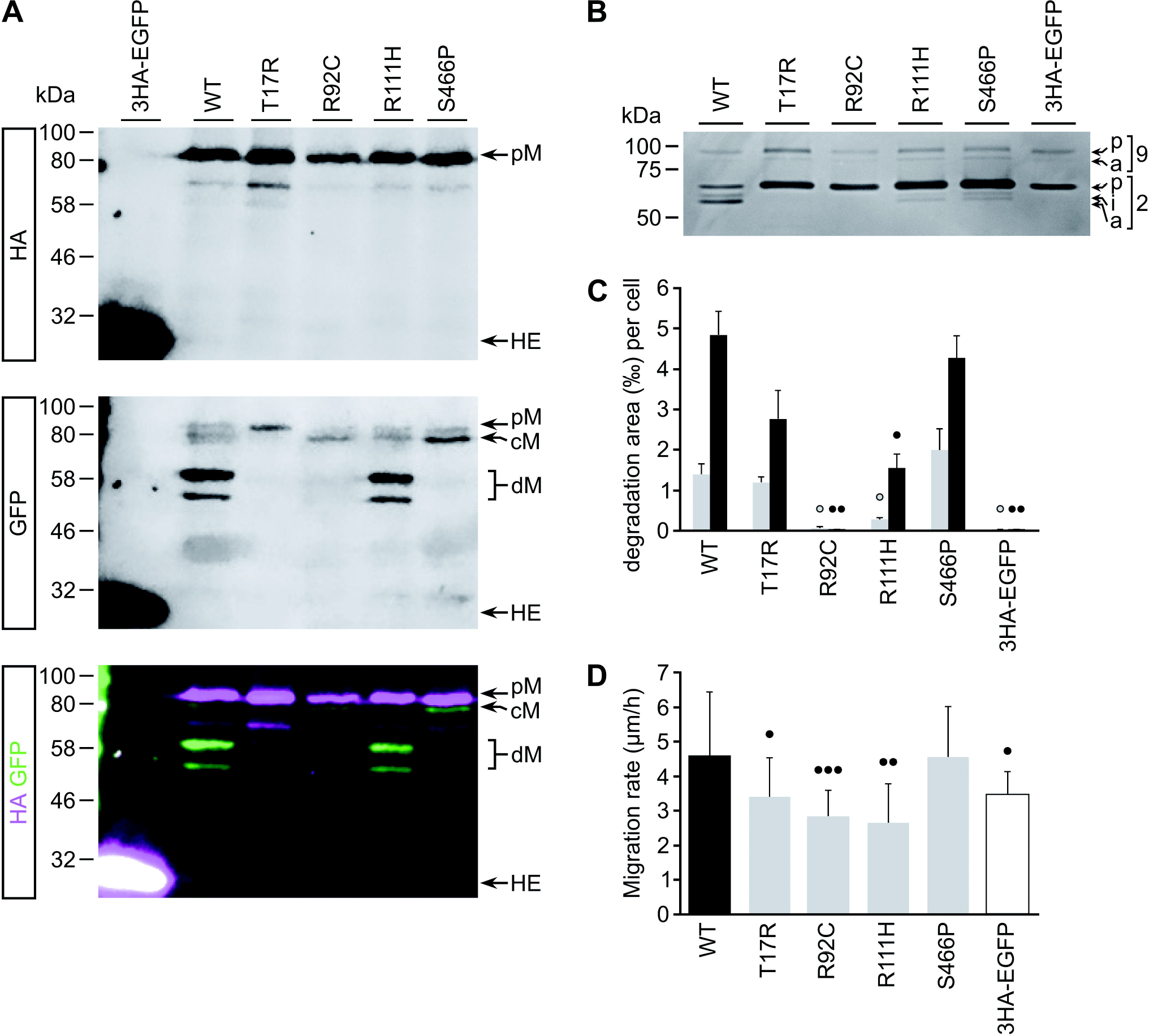
The R111H mutation does not affect MMP14 posttranslational processing, in contrast to mutations T17R, R92C and S466P, yet partially impairs enzymatic and cell migration stimulatory activity. **A**, immunoblot with anti-HA (top panel) or anti-GFP (middle panel) antibodies assessing the posttranslational processing of WT and mutant 3HA-MMP14-EGFP fusion proteins exogenously expressed in MRC5 cells. For all fusion proteins, the full-length pro-protein (pM) can be detected. For all except the T17R mutant, an additional form (cM) can be detected in which the HA tag has been removed. Additional degradation products (dM) can only be detected in MMP14 WT and MMP14 R111H expressing cells. In cells expressing 3HA-EGFP, a strong band corresponding to the fused tags can be detected (HE). **B**, gelatin zymography of media conditioned by MRC5 cells exogenously expressing the WT or mutant 3HA-MMP14-EGFP fusion proteins. Cells expressing 3HA-EGFP control do not activate pro-MMP2 (p2), whereas MMP14 WT expressing cells activate pro-MMP2 to its intermediate (i2) and active (a2) form. The T17R and R92C mutations completely abolish pro-MMP2 activation, whereas the R111H and S466P mutants retain residual activity. Expression of MMP14 fusion proteins does not affect activation of MMP9 (p9 and a9). **C**, average degraded Cy3-gelatin surface area per MMP14 fusion protein expressing MRC5 cell at 4h (grey bars) and 20h (black bars) post seeding. The R92C mutation completely abolishes gelatin degradation, whereas the R111H mutant retains some residual activity. In contrast, the T17R and S466P mutations do not impair gelatin degradation. Error bars indicate SEM of biological triplicates. **D**, average migration rate of MMP14 fusion protein expressing MRC5 cells on fibronectin. Expression of WT MMP14 significantly stimulates migration compared to 3HAEGFP expression, whereas the T17R, R92C and R111H mutations impair this stimulatory effect. In contrast, the S466P mutation does not affect MMP14-dependent migration. Significance levels: dot, p < 0.05; double dot, p < 0.01, triple dot, p < 0.001.

The human mutation p.Thr17Arg (T17R) has previously been reported as causing WS [12]. This substitution is located within the conserved N-terminal signal peptide sequence of MMP14. T17R is thought to prevent proper localization to the ER and subsequent processing of MMP14. Accordingly, whilst the pro-enzyme was observed at the cell membrane by membrane fractionation and surface biotinylation, there was an absence of active MMP14 [12]. We did not observe the T17R mutant at the cell membrane by immunofluorescence (IF, Fig. 1C(ii) and Fig. S4(ii)), most likely due to differences in sensitivity between assays used. Rather, we found MMP14 T17R distributed throughout the cytoplasm. The (partial) co-localization of HA and EGFP tags observed by IF, as well as the overlapping HA and EGFP bands observed by WB (Fig. 2A lane 3) suggest that the T17R mutation impairs MMP14’s signal sequence removal.

To further probe the relationship between MMP14 domains and disease, we generated double-tagged human versions of two MMP14 mutants associated with murine phenotypes that strongly resemble WS. The corresponding mutations are located in distinct MMP14 domains yet cause disorders that are virtually indistinguishable. The p.Arg92Cys (R92C) substitution, underlying the *Sabe* (small and bugged-eyed) phenotype, affects a conserved R^89^-R-P-R-C^93^ furin cleavage motif [17]. As seen in Figures 1C(iii) and S4(iii), MMP14-R92C-EGFP was absent from the cell membrane, suggesting that R92C might impair the prodomain processing that is required for cell surface localization. The removal of the HA tag was confirmed by western blot (Fig. 2A lane 4), however the lack of any additional bands suggests that R92C impairs subsequent processing of MMP14.

The second murine mutation, p.Ser466Pro (S466P), causes the *Cartoon* phenotype [18]. Serine 466 is a highly conserved residue in blade 4 of MMP14’s Hemopexin-like (Hx) domain, which is required for enzyme maturation and trafficking as well as for homodimer interactions [19, 20]. Figure 1C(v) shows extensive perinuclear co-localization of HA and EGFP in cells expressing HAMMP14-S466P-EGFP. Membrane localization of S466P mutant protein (Fig. S4(v)) was markedly reduced compared to WT (and R111H). S466P does not seem to affect the removal of the SP and HA tag (Fig. 2A lane 6), although the reduced intensity of lower bands when compared to those observed for MMP14 WT and R111H suggests that this single amino acid substitution in the Hx domain compromises MMP14 processing.

### MMP14 R111H retains partial pro-MMP2 hydrolyzing activity

Since MMP14 R111H seemed to be processed and trafficked normally, we next assessed the functionality of this mutant with respect to pro-MMP2 activation, utilizing gelatin zymography [7]. First, we determined that medium conditioned by 3HA-EGFP expressing MRC5 cells did not activate pro-MMP2 (Fig. 2B lane 6), consistent with low endogenous MMP14 levels in these cells. Subsequently, we assessed the pro-MMP2 activating potential of media conditioned by cells expressing the tagged WT or mutant MMP14 fusion proteins. As shown in Figure 2B, conditioned media from cells expressing the WT fusion protein converts pro-MMP2 to its intermediate and active forms (lane 1), indicating that the exogenously expressed WT fusion protein is functionally active. Both R111H (Fig. 2B lane 4) and S466P (lane 5) mutants retained partial activity. In contrast, T17R (Fig. 2B lane 2) as well as R92C (lane 3) abrogated pro-MMP2 processing.

### MMP14 R111H retains partial gelatinolytic activity

MMP14 has a key role in mediating invasive cell behavior [7]. Thus, we determined the effect of MMP14 mutations on invasive extracellular matrix (ECM) degradation. We used Millipore’s QCM™ Gelatin Invadopodia Assay system (Cat. No. ECM671) combined with live cell imaging. The results are summarized in Figure 2C, indicating the extent to which MRC5 cells expressing WT or mutant tagged MMP14 were capable of digesting the gelatin matrix on which they were seeded. In line with our previous localization and zymography data, expression of MMP14 WT resulted in significantly more gelatin degradation than expression of 3HA-EGFP. R92C abolished all gelatinase degradation, whereas R111H resulted in a significant reduction yet retained partial activity. Intriguingly, in this assay the S466P mutant retained full activity when compared to the WT protein, despite membrane localization for this mutant being impaired.

### MMP14 R111H has a reduced cell migration stimulatory potential

The gelatin invadopodia assay could not be used to assess MMP14’s ability to promote cellular migration. We repeatedly observed that cells expressing tagged MMP14 seemed to adhere strongly to the poly-L-lysine coated coverslip in areas where gelatin was degraded, such that when the cells finally did move, fragments of cell membrane were left behind (data not shown) resulting in cell death. We hypothesized that abnormal cell migration might contribute to the disease phenotype, as knockdown of *sh3pxd2a* was previously observed to reduce the migrational velocity and podosome formation of zebrafish neural crest cells, resulting in craniofacial malformations [21]. We therefore subjected cells expressing WT or mutant MMP14 fusion proteins to a phagokinetic assay as previously described [22]. Given the cells’ strong adherence to gelatin, we used fibronectin (1 μg/ml) as the substrate. As shown in Figure 2D, the T17R, R92C and R111H mutants demonstrated significantly reduced migratory behavior as compared to MMP14 WT, however without showing a clear correlation with the severity of the associated clinical phenotypes. Unexpectedly, motility of cells expressing MMP14 S466P was not impaired compared to MMP14 WT expressing cells.

### Knockout of mmp14a/b in Danio rerio recapitulates key aspects of the WS phenotype

Our observations suggest that loss of MMP14’s catalytic activity might be a prime determinant of the WS phenotype. To better understand this link, we decided to generate a zebrafish (*Danio rerio*) knockout (KO) model of WS. *D. rerio* is a well-established vertebrate model system for the study of ossification and has several advantages over mice, such as rapid external embryonic development, the ability to generate large numbers of offspring and relatively low cost [23, 24]. *MMP14* is well conserved between humans and zebrafish, consistent with significant functional overlap even though the latter have two copies, *mmp14a* and *mmp14b*, due to genomic duplication [25, 26]. Previous work using Morpholino oligonucleotide (MO)-mediated knockdown suggested that loss of *mmp14a* expression results in defective craniofacial morphogenesis, and hinted at abnormal matrix composition of craniofacial cartilage *Anlagen*. Observations on the consequences of *mmp14b* knockdown are conflicting. It has been proposed that *mmp14a* and *mmp14b* might have a role in cellular migration during gastrulation [25, 27]. However, it was later shown that this was probably a p53-mediated off-target MO effect, although additional non-p53-dependent off-target apoptosis might have contributed as well [28, 29].

In the light of these considerations, and because our data indicate that the mutation in our patients causes a loss of function, using CRISPR/Cas9 we knocked out *mmp14*a and *mmp14b* in zebrafish [30]. Targeting exon 4 in both *mmp14*a and *mmp14b* genes resulted in the introduction of 2bp and 8bp deletions respectively (Fig. S5A), which are predicted to cause a frameshift and premature stop (p.S183Rfs23X and p.F190X respectively). Using qPCR, we showed that the resulting mutant mRNA undergoes nonsense-mediated decay (Fig. S5B & C). We further confirmed that any residual protein product would not contain the WT sequence C-terminal to the mutation site (Fig. S5D).

In subsequent crosses, fish with all possible combinations of WT and mutant *mmp14a/b* alleles were generated. Independent of the parental genotype, *mmp14a*^Δ/Δ^;*mmp14b*^Δ/Δ^ fish (hereafter referred to as *mmp14a/b* KO) developed a gradually worsening phenotype, first noticed around 30 days post fertilization (dpf) and most striking at adult age. Other genotypes were not associated with any overt abnormalities. The double KO phenotype includes decreased total body length (20.6 mm vs. 32.5 mm for WT at 90 dpf, p < 0.0001), a relatively small head showing dorsal hyperextension in most individuals (86% vs. 0%, p < 0.0001), exophthalmos, a short operculum, and thoracic kyphosis (Fig. 3A & B). Although *mmp14a/b* KO fish are born at Mendelian ratio, they have a shortened average life span of about 4.5 months and fail to reproduce. Microcomputed tomography (μCT) at 90 dpf revealed that *mmp14a/b* KO fish have a significantly reduced skull bone mineral density (BMD, 891.5 mg/cc vs. 959.0 mg/cc, p < 0.05) compared to WT fish (Fig. 3C & D), and confirmed Weberian-prehemal hyperkyphosis (angle formed by line connecting center of Weberian vertebral bodies with line connecting prehemal vertebral bodies is 40.7° in *mmp14a/b* KO fish vs. 13.8° in WT, p = 0.0002). Overall, our *mmp14a/b* KO fish recapitulates key aspects of WS and previous murine *Mmp14* mutant and KO models [1, 16–18, 31–33].

**Figure 3.**
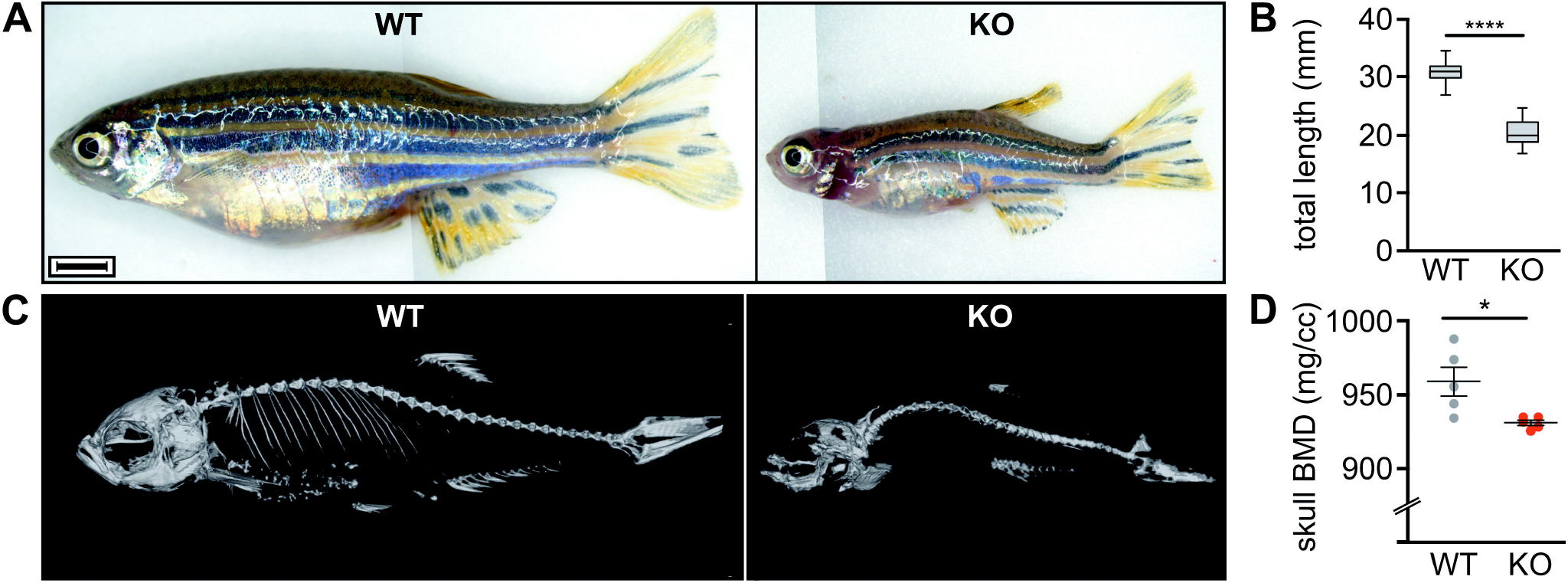
The *mmp14a/b* KO zebrafish recapitulate key aspects of the WS phenotype. **A**, gross anatomy photographs of 3-month-old WT and *mmp14a/b* KO fish of respective average size; lateral view, anterior to the left. The phenotype of *mmp14a/b* KO fish includes a relatively small, up-tilted head with relatively large, protruding eyes and a short operculum. Limited field of view necessitated stitching of multiple photographs together, causing the vertical line in the images shown in panel A. Scale bar equals 2 mm. **B**, at 90 dpf, *mmp14a/b* KO fish have a significantly shorter total body length compared to WT fish (p < 0.0001). A minimum of 21 individuals per genotype was measured. **C**, 3D reconstruction of μCT scans of 3-month-old WT and *mmp14a/b* KO fish; lateral view, anterior to the left. Compared to WT fish, the *mmp14a/b* KO fish have Weberian-prehemal hyperkyphosis. **D**, the *mmp14a/b* KO fish have a reduced skull bone mineral density (BMD, p < 0.05), giving the appearance of missing skeletal elements in the shown 3D reconstruction (C). BMD was assessed for 5 individuals per genotype. The individuals imaged in panel C are different from the ones shown in panel A.

### mmp14a/b KO fish have abnormal endochondral and membranous ossification

Next, we sought to identify the disturbed process(es) underlying the *mmp14a/b* KO skeletal phenotype. At 5 dpf, larval craniofacial cartilage elements are qualitatively normal in size and shape (Fig. S6A & B). Subsequent mineralization of the larval axial skeleton proceeded at a normal pace (Fig. S6C-E). At 30 dpf, the first differences in calvaria shape became apparent (Fig. 4A). Mid-sagittal sections of 90 dpf fish stained with haematoxylin and eosin (H&E) revealed irregular undulation of the membranous ossifying frontal and parietal bones of *mmp14a/b* KO fish, containing cell clusters giving these bones a threadbare appearance (Fig. 4B). The membranous ossifying jawbones contain a relatively small amount of peripheral bone matrix and a disorganized cartilage core (Fig. 4C). Similar abnormalities are seen in the endochondral ossifying supraoccipital bone (SOC), with additional cell-free areas in the cartilage core, devoid of proteoglycans (Fig. 4E) [34]. The relative absence of bone matrix likely accounts for the low skull bone mineral density (BMD) of *mmp14a/b* KO fish. Another striking finding in *mmp14a/b* KO fish is the ventral extension of the SOC and second supraneural (SN2), focally compressing the spinal cord against the first Weberian vertebral body (Fig. 4D). The resulting compromise of spinal cord integrity might contribute to the shortened lifespan of KO fish. This interpretation is supported by observations of an abnormal swimming pattern characterized by tumbling movements, typically observed in KO fish 1-2 days prior to death (Movie S1). Picrosirius red (PSR) staining of additional sections revealed abnormal collagen content of jawbones and the SOC of *mmp14a/b* KO fish, those bones lacking the collagen-rich cortex observed in WT fish (Figs. 4E and S7). Polarized differential interference contrast imaging of PSR stained sections revealed no overt differences in birefringence, suggesting collagen deposition is unaffected by *mmp14a/b* KO (Fig. S7) [35]. Finally, membranous ossifying Weberian vertebral bodies in *mmp14a/b* KO fish were irregularly shaped, with clusters of multinucleated cells in their dorsal aspect (Fig. 4G). Taken together, these findings highlight the importance of Mmp14-dependent collagen remodeling in bone formation.

**Figure 4.**
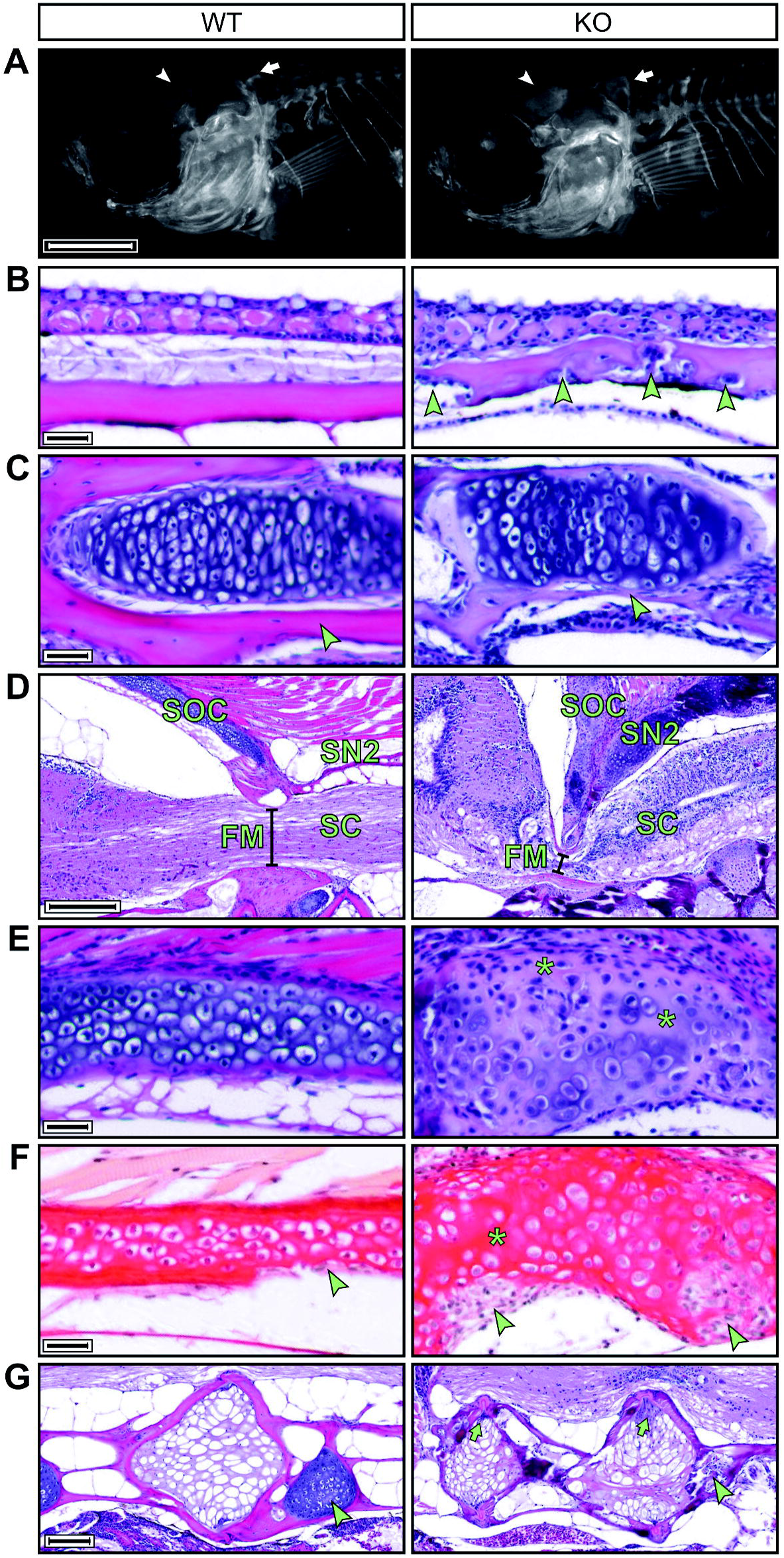
The *mmp14a/b* KO zebrafish have abnormal enchondral and membranous ossifying skull bones and Weberian vertebrae. **A**, fluorescence microscopy images of 30 dpf WT and *mmp14a/b* KO juveniles, whole mount stained for calcified bone with alizarin red; lateral view, anterior to the left. At 30 dpf, the frontal bones (arrowhead) and the supraoccipital bone (SOC, arrow) of *mmp14a/b* KO fish are shaped differently compared to age and size-matched (10.2 mm standard length) WT fish. **B-E**, H&E stained midsaggital sections of 90 dpf fish; anterior to the left except for the *mmp14a/b* KO section shown in panel E, which is rotated (anterior at the bottom) for clearer comparison with the corresponding WT section. At 90 dpf, the frontal bones (B) of *mmp14a/b* KO fish are irregularly thickened and contain cell clusters (arrowheads). The dentary bone (C) and SOC (E) of *mmp14a/b* KO fish contain a relative large amount of disorganized cartilage and small amounts of bone matrix (arrowheads). The SOC additionally shows cell-free areas that have lost basophilia, indicating lack of proteoglycans (E, asterisks). The SOC and second supraneural (SN2) form a sharp angle and are ventrally extended in *mmp14a/b* KO fish, impinging the spinal cord (SC) at the foramen magnum (FM, compare diameter indicated by black line in panel D). **F**, sagittal picrosirius red stained sections of the same fish as shown in panel E (same orientation as in panel E) reveal the SOC of *mmp14a/b* KO fish lacks a collagen-rich peripheral bone matrix, but instead contains large cell-rich areas (arrowheads) as compared to WT fish. In contrast, cell-free regions in the cartilage core of *mmp14a/b* KO fish are relatively intensely stained. **G**, H&E stained midsaggital sections (anterior to the left) demonstrating Weberian vertebral bodies of 90 dpf *mmp14a/b* KO fish are irregularly shaped and contain cell clusters (arrows), while the intervertebral cartilage is absent (arrowhead) compared to WT fish. Scale bar in A equals 1 mm, scale bars in B, C, E and F equal 20 μm, scale bar in D equals 200 μm, scale bar in G equals 100 μm.

## Discussion

Our *in vitro* results, summarized in Table 1, provide evidence that a mild manifestation of WS as observed in our patients is caused by a novel homozygous missense mutation that partially impairs the catalytic activity of MMP14. Our observations suggest that MMP14’s ability to activate pro-MMP2 may be the most important determinant of disease severity in humans, as T17R seemed to affect both gelatinase activity and cell migration to a lesser extent than R111H, whereas it completely abrogated pro-MMP2 activation. The importance of MMP2 is underscored by previous reports that its loss can also cause WS and highly similar phenotypes [11, 36, 37].

**Table 1.** Overview of the effects of studied MMP14 mutations on MMP14 processing and subcellular localization, pro-MMP2 cleavage, gelatin degradation and cell migration.

|  | MMP14 | MMP14 | Pro-MMP2 | Gelatin | Cell |
| --- | --- | --- | --- | --- | --- |
| MMP14 | processing | localization | cleavage | degradation | migration |
| WT | √ | √ | √ | √ | √ |
| T17R | × | × | × | √ | ↓ |
| R92C | × | × | × | × | ↓ |
| R111H | √ | √ | ↓ | ↓ | ↓ |
| S466P | × | ↓ | ↓ | √ | √ |

Tick mark, unaltered; arrow, impaired; cross, severely impaired to absent.

Our findings on MMP14 processing are largely in line with previous work, however some seem contradictory. In contrast to previous observations, we did not observe T17R at the cell membrane. This apparent discrepancy might be due to greater sensitivity of the surface biotinylation assay employed by Evans et al. as compared to our immunofluorescent detection [12]. Our findings on the murine disease-causing R92C mutation seem to contradict previously published observations that the R^89^-R-P-R-C site is shielded from furin, and that the mature enzyme is generated by cleavage at the R^108^-R-K-R-Y motif [13]. Rather, R92C seems to abolish prodomain cleavage. Accordingly, pro-MMP2 activation and gelatinase activity were abrogated and the protein’s ability to support migration impaired.

While our observations on T17R, R92C and R111H are consistent with the prodomain’s known role in MMP14 trafficking and activity, the effects of S466P are less easily explained. As expected from a damaging missense mutation in 4^th^ blade of the Hx domain, it had a pronounced effect on protein processing and trafficking. Yet, the mutant protein showed only reduced activity in gelatin zymography and almost normal activity in the gelatin invadopodia assay. The mutation also did not affect MMP14’s ability to support cellular migration. The most straightforward explanation seems to be that there is sufficient protein still present at the cell surface for the cells to retain near-normal activity in our assays. This is consistent with previously published data showing some membrane localization of MMP14 after its Hx domain had been deleted or swapped out for that of MMP17 [38]. We do not yet know how to reconcile the severe *Cartoon* phenotype, which so strongly resembles that of *Sabe*, or a full *Mmp14* KO, with near normal activity of the S466P mutant. As a possible explanation, we suggest that this mutation might affect functionality of other cell types, such as macrophages and osteoblasts, to a greater extent than that of fibroblasts.

To further assess the effects of loss of MMP14’s catalytic function on the organismal level, we generated a zebrafish *mmp14a/b* KO model. Our *in vivo* results support an important role for MMP14 in collagen remodeling during ossification. Similar to existing mouse models, the *mmp14a/b* KO fish recapitulates essential aspects of the WS skeletal phenotype including a short stature, craniofacial malformations, reduced bone density and abnormal spinal curvature [1, 16, 18, 20, 31–33]. Although the humans, mice and zebrafish all have shortened life spans, the cause of death differs between species. Where WS patients reportedly die from heart failure, the mice succumb to wasting, possibly resulting from malnutrition due to the facial abnormalities [17, 18, 31–33]. Although feeding problems could contribute to the death of *mmp14a/b* KO fish, spinal cord impingement, not observed in humans and mice, seems to be the main cause of death.

In contrast to previous morpholino studies in zebrafish, *mmp14a/b* KO larvae developed normal craniofacial cartilage elements [25, 27]. Subsequent mineralization of the larval skeleton during metamorphosis proceeded normally as well, and the first subtle differences in skull shape only became apparent at juvenile age. The gradually worsening phenotype observed in humans, mice and zebrafish further suggests that loss of MMP14 disrupts later stages of skeletal remodeling. Consistent with our *in vitro* data, these observations argue against a major role for MMP14 supporting cellular migratory and invasive behavior *in vivo*. Although the presence and effect of maternal transcripts in our *mmp14a/b* KO fish cannot be ruled out, the normal skeletal patterning in WS patients and *Mmp14* KO mice supports this notion.

Apart from species-specific differences in bone morphology and ossification mechanisms, the skeletal abnormalities in *mmp14a/b* KO fish are comparable to those observed in *Mmp14* KO mice. Similar to the mice, both endochondral and intramembranous ossification is impaired in *mmp14a/b* KO fish [33]. During these types of ossification, bone forms within, respectively in close association with, a cartilage template [39, 40]. It was shown in *Mmp14* KO mice that these cartilage templates are not properly replaced, respectively removed, due to impaired MMP14-dependent collagen remodeling [32, 33, 41]. This is reflected in the *mmp14a/b* KO fish, where cartilage cores of skull bones are relatively large, disorganized and have altered collagen content, with bone matrix being relative sparse. The cell-filled lacunae in *mmp14a/b* KO fish calvariae are reminiscent of the excessive osteoclastic bone resorption in *Mmp14* KO mouse parietal bone [32–34, 40, 41]. Impairment of additional processes that depend on MMP14’s catalytic activity, including stimulating osteoblast differentiation and activity while having an opposite effect on osteoclasts, might contribute to the *mmp14a/b* KO skeletal phenotype [33, 41–44]. Further studies are needed to assess to which extent these processes underlie the skeletal abnormalities in *mmp14a/b* KO fish.

Taken together, our *in vitro* and *in vivo* studies highlight the significance of impaired proteolytic MMP14 activity underlying the WS skeletal phenotype. This observation has therapeutic implications. Attempts have been made in the past to treat WS and related disorders with bisphosphonates, with limited success [45, 46]. Bisphosphonates exert their therapeutic effect mostly by inhibiting osteoclasts [47]. Whereas our observations as well as previously published work do support a role for osteoclasts in the WS bone phenotype, the major problem seems to be defective collagen remodeling. Thus, we would propose that any treatment of WS should seek to address this fundamental issue. Our zebrafish model could be used to develop and test such treatments, which might also be able to address osteoporosis and other conditions with decreased bone density.

## Materials and methods

### Cell culture

MRC-5V1 immortalized human fetal lung fibroblasts were provided by Prof. Alan Lehmann (University of Sussex, Brighton, UK). HT-1080 fibrosarcoma cells were provided by Dr. John Eykelenboom (University of Dundee, Dundee, UK). Cells were grown in 2D culture in high glucose Dulbecco’s Modified Eagle Medium (DMEM, GE Healthcare Life Sciences, Pittsburgh, Pennsylvania; SH3024.01), containing 10% (*v/v*) fetal bovine serum (FBS; GE Healthcare Life Sciences, Pittsburgh, Pennsylvania; A15-101), 100 U/mL penicillin and 100 μg/mL streptomycin (Thermo Fischer Scientific Inc., Waltham, Massachusetts, USA; 15140122) at 37 °C in 100% humidity and 5% CO_2_. Cells were kept growing in log phase and passaged when reaching 70-85% confluence by detaching cells with 0.25% trypsin-EDTA (Thermo Fischer Scientific Inc., Waltham, Massachusetts; 25200-056).

### Cloning and mutagenesis

A series of expression vectors was created encoding 3HA and EGFP double-tagged versions of either WT or mutant human MMP14. The MMP14 WT vector was generated by amplifying *MMP14* (RefSeq NP_004986.1) from total cDNA obtained from MRC-5V1 cells (see Table S1 for primers used) using KOD high fidelity polymerase (Merck) and cloning the amplicon into the pJET3.1 vector (Thermo Fisher). *MMP14* WT cDNA was cloned into the pQCXIB (w297-1, Addgene plasmid #22800) backbone (a gift from Dr. Eric Campeau) with addition of 3HA and EGFP tags in subsequent cloning steps. The WS R111H mutant vector was generated by site-directed mutagenesis (SDM) of codon 111 of the *MMP14* cDNA sequence in the WT vector from CGC into CAC using partially overlapping primers following the protocol of Werler et al. [48]. Likewise, the *Cartoon* S466P mutant vector was generated by SDM of codon 466 from TCA into CCA. The WS T17R mutant vector was created by SDM of codon 17 from ACG into AGG with fully overlapping primers (see Table S1). Likewise, the *Sabe* R92C mutant vector was created by mutating codon 92 from CGA into TGC with fully overlapping primers. The 3HA-EGFP control vector was generated by deletion of the *MMP14* coding sequence from the WT vector by inverse-PCR with fully overlapping primers binding to 24 bp of each linker sequence.

Four expression vectors encoding either WT or mutant *mmp14a* or *mmp14b* with a C-terminal HA tag were generated. cDNA was obtained from 2-month-old WT and *mmp14a*^Δ/Δ^;*mmp14b*^Δ/Δ^ fish, respectively, as described below and respectively WT or mutant *mmp14a/b* coding sequence PCR amplified with the primers listed in Table S2. The forward primers started with zebrafish-optimized kozak sequence 5’-CCAACC-3’, the reverse primer ended with the HA coding sequence. The amplicon was gel-extracted and cloned into the pCR2.1 TOPO TA vector (Thermo; 450641) according to the manufacturer’s protocol. The kozak-*mmp14a/b*-HA insert was subsequently cloned into the pCS2+ backbone.

### Immunofluorescence microscopy

Cells were seeded on No. 1.5H microscope cover glasses (Paul Marienfeld GmbH & Co. KG, Lauda-Königshofen, Germany; 0117580) in 12-well cell culture plates (Corning Inc., Corning, New York, USA; 353043) at a density of 50,000 cells per well. Cells were transfected with 1.2 μg vector DNA using jetPRIME^®^ (Polyplus-transfection^®^ SA, Illkirch, France; 114-15) 24h post seeding and media refreshed 4h post transfection. Non-transfected cells were seeded 18h prior to fixation. Cells were fixed with 4% paraformaldehyde (PFA) in PBS 1x (Santa Cruz Biotechnology, Inc., Dallas, Texas, USA; sc-281692), permeabilized with 0.2% Triton^®^ X-100 (Promega, Madison, Wisconsin, USA; H5141) in PBS 1x (GE Healthcare Life Sciences, Pittsburgh, Pennsylvania; SH30028.02) and blocked with Image-iT^®^ FX Signal Enhancer (Thermo Fisher Scientific Inc., Waltham, Massachusetts, USA; R37107). Cells were incubated with primary antibodies (See Table S3 for antibody details and dilutions) overnight at 4 °C, and with Alexa-fluorophore labelled secondary antibodies (1:200) and phalloidin (1:100) (Thermo Fisher) for 1 h at RT, all dissolved in 3% (*w/v*) bovine serum albumin (BSA; MP Biomedicals, Santa Ana, California, USA; 0219989880) in PBS 1x. Cells were mounted on microscope glasses (Biomedia, Singapore, Singapore; BMH.880102) using Vectashield antifade mounting medium containing 4′,6-diamidino-2-phenylindole (DAPI) (Vector Laboratories, Burlingame, California, USA; H-1200). Z-stacks of cells were recorded using an Olympus IX81 microscope with FV1000 scan head (Olympus Corporation, Tokyo, Japan). Images of randomly selected cells with average signal intensity were taken with a 63X 1.45 NA oil immersion Plan Apochromat objective (Olympus), with fixed laser power and exposure time per experiment. The investigator taking and analyzing images was blinded for the transfection of the cells. Raw images were processed with Fiji software (U. S. National Institutes of Health, Bethesda, Maryland, USA; ImageJ version 2.0.0-rc-15/1.49k, adapted by IMU).

### Western blotting

For endogenous MMP14 and analysis of double-tagged MMP14, cells were seeded at a density of 100,000 cells per well of a 6-well plate. Cells were either transfected with pQCXIB vectors as described above or left non-transfected. Cells were harvested using trypsin-EDTA and whole cell lysate (WCL) was obtained using NP-40 lysis buffer (150 mM NaCl, 1% NP-40, 250 mM Tris pH 7.3) supplemented with protease and phosphatase inhibitors (Roche). Protein concentration was determined by Bradford protein assay (Sigma). An appropriate volume of Laemmli sample buffer was added to equal amounts of total protein lysate followed by boiling for 5 min. Samples were subjected to SDS–PAGE electrophoresis before transfer to an Amersham Hybond™-PVDF membrane (GE Healthcare Life Sciences, Buckinghamshire, UK). Membranes were blocked in 5% (*w/v*) milk (Marvel dried skimmed milk) in 0.1% (*v/v*) Tween-20 (Melford; P1362) in TBS 1x (Sigma; T5912-1L). After overnight incubation at 4°C in primary antibody (see Table S3) appropriately diluted in 0.5% BSA in TBS block buffer, membranes were incubated with secondary IRDye^®^ 680RD donkey anti-rabbit (LI-COR Biosciences, Lincoln, Nebraska, USA; 926-68073) and Alexa fluor 680 donkey anti-mouse (Thermo; A10038) antibody appropriately diluted in block buffer. Blots were imaged on the Odyssey^®^ Fc Imaging System (LI-COR).

For analysis of the effect of *mmp14a/b* frameshift mutations on putative protein products, 400,000 MRC-5V1 cells were seeded in 6 cm dishes. Cells were transfected 24h post seeding with 8 μg pCS2+ vector or left non-transfected using Invitrogen^™^ Lipofectamine 3000 (Thermo; L3000008) according to the manufacturer’s protocol. Media was refreshed 6h post transfection. Cells were harvested 24h post transfection using trypsin-EDTA and WCL obtained using Triton^®^ X-100 lysis buffer (50 mM Tris-HCl pH 7.5, 150 mM NaCl, 1% (*v/v*) Triton^®^ X-100, 10% (*v/v*) glycerol, 0.1% (*w/v*) deoxycholate, 25 mM β-glycerophosphate) supplemented with protease inhibitors (Roche) and phosphatase inhibitors (10 mM NaF and 1 mM Na_3_VO_4_). Protein concentration was determined by Bradford protein assay. Samples were mixed with Laemmli sample buffer and boiled as described above. Samples were subjected to SDS–PAGE electrophoresis before transfer to a PVDF membrane (Merck; ISEQ00010). Membranes were blocked in 5% (w/v) milk (Biorad; 170-6404) in 0.1% (*v/v*) Tween-20 in TBS 1x and incubated with rabbit anti-HA (Cell Signaling Technology Inc., Danvers, Massachusetts, USA; 3724) or rabbit anti-β-actin (Cell Signaling; 4967) primary antibodies at 1:1000 or 1:4000, respectively, in 5% (*w/v*) milk in 0.1% (v/v) Tween-20 in TBS 1x. Membranes were incubated with horseradish peroxidase (HRP)-conjugated goat anti-rabbit secondary antibody (1:3,000) and Streptactin (1:10,000) in 4% (*w/v*) milk in 0.1% (v/v) Tween-20 in TBS 1x. Protein bands were visualized by ECL Western Blotting substrate mix (Thermo; 32106) on the BioRad Chemidoc^™^ and gel images edited in Image LabTM 5.1 build 8 software (Biorad).

### Zymography

Gelatin zymography was performed as previously described [49]. Briefly, MRC-5V1 cells were transfected with the aforementioned pQCXIB vectors. Culture medium was changed to FBS-free DMEM 24h post transfection to allow conditioning for 24h, after which conditioned medium was harvested and mixed with equal volumes of 2x sample buffer (0.125 M Tris-HCL, pH 6.8, 20% (*v/v*) glycerol, 4% (*w/v*) SDS, 0.005% (*w/v*) bromophenol blue (Sigma; B-6131)). Samples were subjected to SDS-PAGE electrophoresis in 10% resolving gel containing 0.1% gelatin. After electrophoresis, proteins were renatured in freshly made 1X Zymogram Renaturing Buffer (Triton X-100 2.5% (*v/v*) in ddH_2_O) for 30 min at RT. The gel was subsequently incubated in 1x Zymogram Developing Buffer (0.01 M Tris-HCL pH 7.5 (Sigma; 10812846001), 1.25% (*v/v*) Triton X-100, 5mM CaCl_2_ (Kanto Chemical Co., 07058-00)) for 30 min at RT to allow equilibration. Developing buffer was refreshed and the gel was incubated at 37 °C overnight.

Finally, the gel was stained with Coomassie Blue R-250 (Thermo; 20278) 0.5% (*w/v*) for 30 min, and subsequently washed in destaining solution (water: methanol acetic acid at 5: 4: 1) for 4 times 15 min. The gel was imaged with the Bio-Rad Image Lab system and the image analyzed in Fiji software.

### Gelatin digestion assay

The QCM^™^ Gelatin Invadopodia Assay system (Merck Millipore, Billerica, Massachusetts, USA; ECM671) was used according to the manufacturer’s protocol. In brief, a 4-chamber glass bottom dish (Cellvis, Mountain View, California, USA; D35C4-20-1.5-N) was coated with poly-L-Lysine, glutaraldehyde and Cy3-labelled gelatin in sequence. The coated dish was sterilized with 70% ethanol for 30 min, followed by DMEM quenching for 30 min at RT. MRC-5V1 cells were transfected with the appropriate pQCXIB vectors 24h prior to seeding onto the coated chamber dish. The ability of the transfected cells to digest the Cy3-labelled gelatin was determined at 3h and 20h post seeding using an EVOS fluorescence microscope (Thermo; AMF4300). Average degradation per green fluorescent cell (degradation factor, d) was calculated by automated image analysis in Fiji and assessed for statistical difference by a 2-sided student’s T-test in Microsoft^®^ Excel^®^ (Microsoft Corporation, Redmond, Washington, USA; version 14.6.6 (160626)). To quantify the number of transfected cells (N) in a given field, a find-object macro was created in Fiji. Threshold was manually adjusted to cover the entire degradation area (A) that was measured. The degradation factor of each MMP14 mutant was defined as d = A/N. Three independent experiments (N≥103) were carried out. For qualitative analysis, fluorescent cells were identified 2h post seeding and z-stacks recorded ever 15 min for 5h with a spinning disk (Yokogawa, Tokyo, Japan; CSU-22) confocal microscope (Nikon, Tokyo, Japan; Nikon Ti inverted) equipped with the Nikon Perfect Focus System. A maximum intensity projection was performed in MetaMorph 7.8.8.0 software (Molecular Devices, Sunnyvale, California, USA) at the end of the experiment.

### Phagokinetic assay

Four-chamber glass bottom dishes were coated with 1 μg/mL Rhodamine Fibronectin (Cytoskeleton, Denver, Colorado; FNR01) according to the manufacturer’s protocol. MRC-5V1 cells were transfected with the aforementioned pQCXIB vectors 24h prior to seeding onto the coated chamber dish. Migration of the transfected cells was recorded by time-lapse imaging starting 2h post seeding with a spinning disk confocal microscope. A z-stack consisting of 16 steps with an interval of 1 μm between successive steps of each field was acquired and a maximum intensity projection was performed in MetaMorph software. Time-lapse images were then subjected to customize cell-tracking analysis in Imaris Image Analysis Software 8.4.1 (Bitplane, Belfast, UK).

### Zebrafish husbandry

*D. rerio* larvae of the AB strain were grown in E3 medium (0.1 mM NaCl, 3.4 μM HCl, 6.6 μM CaCl_2_, 6.6 μM MgSO_4_, pH 7.4) at 28.5 °C at a rearing density of 80-100 individuals per liter until 14 dpf, after which larvae were transferred to a closed water system at a density of 13-16 individuals per litre. Feeding schedule was fixed according to the age of the fish; light/dark (14h/10h) circadian rhythm remained fixed from 5 dpf onwards. From 4 weeks of age onwards, fish were kept at 27.5 °C. All zebrafish experiments were conducted under IACUC licence 140924.

### Genomic editing of mmp14a and mmp14b by CRISPR/Cas9

Optimal CRISPR genomic target sites in *mmp14a* (RefSeq NP_919397.1) and *mmp14b* (RefSeq NP_919395.1) were identified by ZiFiT Targeter Version 4.2 (Zinc Finger Consortium, URL: zifit.partners.org/ZiFiT), ordered as gBlocks^®^ Gene Fragments (Integrated DNA Technologies, Coralville, Iowa, USA), PCR amplified and transcribed into gRNA with the Invitrogen^™^ MEGAshortscript T7 kit (Thermo, Fischer Scientific Inc., Waltham, Massachusetts, USA; AM1354) according to the manufacturer’s protocol [50, 51]. *Cas9* RNA was generated by transcribing *NotI*-linearized pCS2-*nls-zCas9-nls* vector (Addgene, Cambridge, Massachusetts, USA; 47929) with the Invitrogen^™^ mMESSAGE mMACHINETM SP6 Transcription Kit (Thermo; AM1340) according to the manufacturer’s protocol [52]. *D. rerio* zygotes were micro-injected in the yolk cell with ~2.5 nL 1x Danieau’s solution (58 mM NaCl, 0.7 mM KCl, 0.4 mM MgSO_4_, 0.6 mM Ca(NO_3_)_2_, 5.0 mM HEPES, pH 7.6) / 1x PhenolRed (Sigma; P0290) in dH_2_O containing 0.375 ng *Cas9* RNA and 0.375 ng *mmp14a* gRNA or 1 ng *mmp4b* gRNA as previously described [53]. Genomic editing was assessed by lysis of 24 hpf embryos or juvenile/adult fin-clips (in 10 mM Tris, 50 mM KCl, 0.3% (*v/v*) Tween-20, 0.3% (*v/v*) NP-40, and 0.488 mg/mL proteinase K in dH_2_O; pH 8.3) and direct Sanger sequencing of a 200-500 bp gDNA amplicon encompassing the target site using Bigdye Terminators version 3.1 (Thermo; 4337455) and a 3730XL sequencer (Thermo). Chromatography reads were analyzed for frameshift mutations with Poly Peak Parser web tool (Yost lab, Salt Lake City, Utah, USA) and checked manually in SnapGene version 3.3.4 (Clontech Laboratories, Mountain View, California, USA) [54]. Three-month-old F0 mosaic mutants were intercrossed and their F1 offspring genotyped at 2 months of age to identify fish with heterozygous frameshift mutations in *mmp14a* or *mmp14b*. F1 *mmp14a*^+/Δ^ fish were outcrossed with F0 *mmp14b* founders to generate F2 *mmp14a*^+/Δ^;*mmp14b*^+/Δ^ fish, that were subsequently intercrossed to generate all possible *mmp14a/b* genotypes in the F3 generation.

### qPCR verification of mmp14a/b knockout

Twenty 1-5 dpf larvae per genotype were pooled per time point, homogenized in Trizol (Thermo; 15596018) and total mRNA was chloroform/isopropanol precipitated and rinsed with ice-cold 75% (*v/v*) enthanol in diethyl pyrocarbonate (DEPC) treated water (Sigma; 159220). mRNA was resuspended in RNase-free water, DNase treated (Thermo; AM2238) and cleaned-up with the RNeasy Mini Kit (Qiagen; 74106). cDNA was synthesized with the High-Capacity cDNA Reverse Transcription Kit (Thermo; 4368813) and qPCR was performed with SYBR^™^ Select Master Mix (Thermo; 4472919), all according to the manufacturers’ protocols. Primers listed in Table S4 were tested at different concentrations (1.95-500 nM) for qPCR on cDNA of 24 hpf WT embryos in technical triplicate with SYBR^™^ Select Master Mix according to the manufacturer’s protocol. Generation of a single amplicon verified by direct Sanger sequencing. Expression of *mmp14a/b* was normalized to β-actin (and expressed relative to the expression level at 1 dpf for larvae).

### Gross anatomy of larvae, juveniles and adult zebrafish

Fish were sedated with 200 μg/mL ethyl 3-aminobenzoate methanesulfonate (Tricaine/MS222; Sigma; A5040) in system water buffered to pH 7.0-7.5 with NaHCO_3_ (Sigma; S5761) and imaged on moist filter paper with an MZ16 FA fluorescence stereomicroscope (Leica, Wetzlar, Germany) and DFC 300 FX Digital Color Camera (Leica) at 1.4 MPixel resolution at 7.11x and 14x magnification [55]. Images were stitched together in Pixelmator 3.5 Canyon software (Pixelmator Team, Vilnius, Lithuania). Total body length was measured using Fiji software and differences in mean total body length between genotypes was tested for significance by two-sided Student’s t-test. The number of fish with dorsal tilting of the head was analyzed by Fisher Exact test in SPSS software version 22 (IBM, Armonk, New York, USA).

### Microcomputed tomography

Three-month-old fish were euthanized by hypothermic shock and fixed in 4% PFA and dehydrated through graded water into ethanol. Average sized fish were fixed per genotype, except for *mmp14a*^Δ/Δ^;*mmp14b*^Δ/Δ^ for which larger individuals were selected to better match the size of other genotypes. Microcomputed tomography images were acquired using an Inveon CT (Siemens AG, Berlin, Germany) at 55 kVp / 110 mA. The exposure time per projection was 2,500 ms and a binning factor of 2 was used, resulting in a reconstructed pixel size of 35 μm. Planar images were acquired from 181 projections over 360° of rotation in step-and-shoot mode. The images were reconstructed using a Feldkamp cone-beam algorithm. Three-dimensional renders of the skeleton were made with AMIRA software (FEI, Mérignac Cedex, France) with constant window settings. Images were exported as TIFF files and extracorporeal skeletal elements of previously imaged fish were manually removed with Pixelmator software. Raw data was viewed with AMIDE-bin 1.0.5 software (Andreas Loening), individual virtual sections exported as TIFF files and angles measured in Fiji software. Differences in mean bone density and angle of kyphosis between genotypes was tested for significance by two-sided Student’s t-test.

### Histology

Three-month-old fish were euthanized and fixed as described above. After trimming, the fish were placed into cassettes and processed with Sakura VIP Tissue Processor. Fish were dehydrated through graded ethanol into xylene (Chemtech Trading) before paraffin (Leica; 39601006) infiltration. After processing, tissues were embedded into paraffin blocks and sectioned midsaggitally with a rotary microtome into 5 μm thick sections. The slides with sections were dried and placed into an incubator (60°C, 15 min). The sections were deparaffinized and rehydrated through graded ethanol into water. Rehydrated sections were subjected to haematoxylin solution (Richard-Allan ScientificTM; 7231), Bluing (Richard-Allan ScientificTM; 7301), Clarifier (Richard-Allan ScientificTM; 7442) and eosin-phloxine B Solution (AMPL). A separate batch of 5 μm thick sections of all the samples were stained with Weigert’s hematoxylin, followed by Picrosirius red 0.1% (*w/v*) staining and rinsing with two changes of acidified (0.5%) water. Stained sections were dehydrated through graded ethanol into xylene and a cover slip added. Slides were imaged with an automated slide scanner equipped with Zeiss AxioImager Z.2 body, MetaSystems stage, Zeiss Plan-Neofluar 20x/0.5 Ph2 lens, SSCOPED TL light source, CoolCube 1 camera with 1.4 Mpixel resolution controlled by Metafer4 software. Images were viewed with VSViewer V2.1.103 software (MetaSystems GmbH, Altlussheim, Germany).

### Whole mount cartilage and calcified bone staining

Larvae were stained for cartilage and calcified bone according to a protocol adapted from Walker et al. [56]. Larvae were culled by overdose of Tricaine, fixed in 4% PFA for 2h at RT and dehydrated in 50% (*v/v*) ethanol. Larvae were stained with either 0.02% (*w/v*) Alcian blue 8 GX (Sigma; A5268) / 60 mM MgCl_2_ or 40 μg/mL 3,4-dihydroxy-9,10-dioxo-2-anthracenesulfonic acid sodium salt (Alizarin Red, Sigma; A5533) in 70% (*v/v*) ethanol for 14h at RT. Larvae were rehydrated in 50% ethanol (*v/v*). Thirty dpf alizarin red stained larvae were de-stained in 1% (*w/v*) KOH for 45 min at RT. All larvae were bleached with 1.5% (*v/v*) H_2_O_2_ / 1% (*w/v*) KOH for 20 min at RT. Larvae were cleared by going through successive stages (20-50% *v/v*) of glycerol. Fourteen dpf alizarin red stained larvae and picrosirius red stained sections were imaged with a Zeiss AxioImager M2 upright fluorescence microscope with X-Cite^®^ 120Q (120 W) light source (Excelitas Technologies Corp., Waltham, Massachusetts, USA), dsRed filter, DIC polarizer and Zeiss Plan-Neofluar 5X / 0.16 NA, 10X / 0.3 NA and 20X / 0.5 NA lenses. The system was operated with AxioVision version 4.8.2 SP3 software (Zeiss) and images were taken with an AxioCam HRc camera (Zeiss) with 1.4 Mpixel resolution. Alcian Blue stained larvae and 21 and 30 dpf alizarin red stained larvae were imaged with a Leica MZ16FA fluorescence stereomicroscope, equipped with the Leica CLS150 (for brightfield) and MZ16FA (for fluorescence) light sources, dsRed filter and Planapo 1.0x lens (Leica; 10447157). The system was operated with Leica Application Suite software version 2.5.0 R1 (Build 975) and images were taken with a Leica DFC 300 FX R2 camera with 1.4 Mpixel resolution. Differences in mean standard length and vertebral ossification between genotypes was tested for significance by two-sided Student’s t-test.

## Acknowledgments

This work was supported by the Skin Research Institute of Singapore [to M.v.S. and E.Y.T], the Biomedical Research Council Singapore [to M.v.S.], the Agency for Science, Technology and Research [A*STAR Research Attachment Programme to I.d.V.], the Wellcome Trust [DGEM to M.v.S. and B.J.C.], Tenovus Scotland [T15/22 and T15/62 to M.v.S. and B.J.C.], the Victorian Government’s Operational Infrastructure Support Program, and the Australian Government [NHMRC Senior Research Fellowship 110297 and NHMRC Program Grant 1054618 to M.B., NHMRC Career Development Fellowship GNT1032364 to P.J.L.]. The authors acknowledge the invaluable assistance of Dr. Graham D. Wright, Dr. Jaron Liu and Dr. Shuping Lin (Institute of Medical Biology, Microscopy Unit), Dr. Monique N.H. Luijten (Lee Kong Chian School of Medicine, Nanyang Technological University), and Dr. Chee Bing Ong (Advanced Molecular Pathology Laboratory).

## Conflicts of Interest Statement

The authors report no conflict of interest. The authors alone are responsible for the content and writing of this article.

## Abbreviations

BMD: bone mineral density
μCT: microcomputed tomography
DAPI: 4′,6-diamidino-2-phenylindole
DEPC: diethyl pyrocarbonate
DMEM: Dulbecco’s Modified Eagle Medium
ECM: extracellular matrix
EGFP: enhanced green fluorescent protein
FTHS: Frank-Ter Haar syndrome
HA: hemaglutinin
H&E: haematoxylin and eosin
HRP: horseradish peroxidase
Hx: hemopexin
KO: knockout
MMP2: matrix metalloproteinase 2
MMP14: matrix metalloprotease 14
MO: Morpholino oligonucleotide
MONA: multicentric osteolysis, nodulosis and arthropathy
PSR: picrosirius red
*Sabe*: small and bugged-eyed
SDM: site-directed mutagenesis
SH3PXD2B: SH3 and Phox-homology (PX) Domain-containing Protein 2B
SN2: supraneural 2
SOC: supraoccipital bone
WB: Western blot
WS: Winchester syndrome
WT: wild type

